# Calmodulin mutation N54I causes autosomal dominant CPVT in mice

**DOI:** 10.64898/2026.09.12.751164

**Authors:** Daniel J. Blackwell, Nieves Gomez-Hurtado, Kyungsoo Kim, Paxton A. Ritschel, Carlos Tellet-Cabiya, Huan He, Jose R. Pinto, Eric Delpire, Bjorn C. Knollmann

**Affiliations:** Department of Medicine, Vanderbilt University Medical Center, Nashville 37235, Tennessee, United States; Translational Science Laboratory, College of Medicine, Florida State University, Tallahassee, FL; Institute of Molecular Biophysics, Florida State University, Tallahassee, FL, USA; Department of Biomedical Sciences, College of Medicine, Florida State University, Tallahassee, FL, USA; Department of Anesthesiology, Vanderbilt University School of Medicine, Nashville, TN, USA

**Author notes:** **Corresponding Author:** Björn C. Knollmann, MD, PhD, Medical Research Building IV, Rm. 1265, 2215B Garland Ave, Nashville, TN 37232-0575.

**Keywords:** Calmodulin, Catecholaminergic Polymorphic Ventricular Tachycardia, Long QT Syndrome, Calcium, RyR2 Calcium Release Channel, Sarcoplasmic Reticulum

## Abstract

Catecholaminergic polymorphic ventricular tachycardia (CPVT) is an inherited arrhythmia syndrome characterized by stress- or catecholamine-induced ventricular arrhythmias in the absence of overt structural heart disease. Mutations in *RYR2* and *CASQ2* account for most genetically defined cases, although pathogenic variants in the three genes encoding calmodulin (*CALM1–3*) have also been linked to CPVT. Because all three *CALM* genes encode an identical calmodulin protein, pathogenic calmodulin variants are expected to be expressed in only a small fraction of total cellular calmodulin, raising the question of whether this limited abundance is sufficient to produce an arrhythmogenic phenotype in vivo. We generated a heterozygous mouse model carrying the human disease-associated N54I-equivalent mutation, N54I, in *Calm1*. Mutant calmodulin accounted for 13.7% of total cardiac calmodulin, consistent with expression from one of six *Calm* alleles. Under basal conditions, N54I/+ mice exhibited normal growth, survival, cardiac morphology, and surface electrocardiogram parameters. However, cardiomyocytes isolated from N54I/+ mice had increased rates of RyR2-mediated spontaneous calcium release. Following catecholaminergic challenge with isoproterenol and caffeine, N54I/+ mice exhibited significantly more premature ventricular contractions and arrhythmias than wild-type littermates. Exercise challenge in conscious mice similarly provoked ventricular ectopy and ventricular tachycardia. In addition, N54I/+ mice exhibited abnormalities of atrial and sinoatrial electrical activity, including premature atrial contractions, ectopic P waves, atrioventricular conduction slowing, and beat-to-beat variability. These findings demonstrate that expression of the N54I calmodulin variant from a single *Calm1* allele is sufficient to produce a CPVT phenotype in vivo. This model provides experimental evidence linking a human disease-associated calmodulin variant to arrhythmogenesis and demonstrates the functional dominance of mutant calmodulin in the heart.

**One Sentence Summary:** A single CALM1 N54I allele causes catecholaminergic polymorphic ventricular tachycardia in mice

## Introduction

The calcium-modulated protein (calmodulin, CaM) is a ubiquitous messenger protein that regulates many cellular processes, including inflammation, metabolism, smooth muscle contraction, memory potentiation, and gene expression, among others.^1,2^ In the heart, CaM regulates several ion channels on the sarcolemma via direct interaction or via activation of the Ca/CaM kinase II (CaMKII).^3^ Within the sarcoplasmic reticulum (SR), CaM regulates calcium release both via CaMKII phosphorylation and via direct interaction with the cardiac ryanodine receptor (RyR2). It also regulates SR calcium uptake via CaMKII phosphorylation of phospholamban.^3^ Pathogenic mutations in CaM may alter its calcium binding or affinity for direct protein interactions and have been documented in several genetic arrhythmia syndromes including catecholaminergic polymorphic ventricular tachycardia (CPVT),^4^ long QT syndrome (LQTS), and idiopathic ventricular tachycardia.^5^

The importance of CaM is underscored by the unusual genetic organization of the human *CALM* genes: three independent genes, *CALM1, CALM2*, and *CALM3*, encode the same CaM protein sequence, and all three *CALM* genes are expressed in the heart.^6^ Consequently, a heterozygous mutation in a single *CALM* gene is expected to contribute only a fraction of the total cardiomyocyte CaM pool. Nevertheless, pathogenic variants in all three *CALM* genes have been reported to cause autosomal-dominant cardiac arrhythmia syndromes.^7^

CPVT is characterized by catecholamine- or stress-induced ventricular arrhythmias caused by aberrant SR calcium release. Untimely calcium release causes the electrogenic sodium-calcium exchanger to remove calcium from the cell and drive sodium into the cell; this reverse mode transport can produce delayed after-depolarizations.^8^ These delayed after-depolarizations may generate ectopic beats when located in subendocardial cardiomyocytes near Purkinje fibers,^9^ which can lead to fatal arrhythmias. CPVT typically manifests in adolescence and has a high mortality rate if left untreated.^10^

Mutations in *RYR2*, which encodes the cardiac SR Ca^2+^ release channel, account for approximately half of dominantly inherited CPVT, whereas recessive CPVT is commonly associated with mutations in *CASQ2*. Experimental mouse models of CPVT have established a direct relationship between abnormal SR Ca^2+^ release and catecholamine-induced ventricular arrhythmias. For example, *Casq2* knockout mice exhibit increased diastolic SR Ca^2+^ leak, spontaneous Ca^2+^ release, triggered activity, and stress-induced ventricular arrhythmias despite relatively preserved cardiac structure and contractile function under basal conditions.^11^

More recently, calmodulin mutations have been discovered in patients with CPVT. The first disease-associated *CALM1* variant (N54I) was identified in a Swedish family with an autosomal dominant CPVT-like syndrome and was initially reported as p.Asn53Ile in humans.^4^ The same report identified a second de novo *CALM1* variant (N98S), p.Asn97Ser, in another individual with clinical features of CPVT.^4^

The clinical identification of CALM variants in clinical cases of CPVT raises an important mechanistic question: how can a mutation expressed from only one of six *CALM* alleles produce a dominant arrhythmia phenotype? RyR2 exists as a homotetrameric Ca^2+^ release channel, with CaM binding to multiple sites on the channel complex. Thus, relatively small alterations in the functional properties of mutant CaM could have disproportionate effects on SR Ca^2+^ release if mutant CaM interacts with and alters the gating of RyR2 channels. CaM is a direct regulator of RyR2, and CaM binding influences RyR2 channel gating. Our previous biochemical and in vitro studies utilizing recombinant wild-type and mutant CaM proteins associated with CPVT and LQT demonstrated that unlike LQTS-associated CaM variants,^6^ recombinant CaM-N54I has normal calcium binding affinity.^12^ However, CaM mutations that produce a CPVT phenotype in humans (CPVT-CaMs) bind to RyR2 with higher affinity than wild-type CaM and activate rather than inhibit RyR2 channels.^12^ Consequently, RyR2-mediated SR calcium leak is enhanced in cardiomyocytes. Importantly, a mixture containing only one part mutant CaM N54I to eight parts wild-type CaM was sufficient to increase spontaneous Ca^2+^ release,^12^ providing a potential molecular explanation for the dominant inheritance of CPVT-associated CaM mutations.

Despite this mechanistic evidence from in vitro studies with recombinant CaM-N54I, it remains unclear whether expression of a CPVT-associated CaM mutation from a single endogenous allele is sufficient to reproduce the arrhythmia phenotype in vivo, especially since N54I has been labeled as a likely benign variant on a functional analysis of mutant CaMs in a deep-mutational scan.^13^ We therefore generated a knock-in mouse carrying the N54I mutation in *Calm1*. We sought to determine the abundance of mutant CaM, and whether this was sufficient to reproduce a CPVT phenotype in vivo, characterize the associated cellular Ca^2+^ phenotype, and determine whether the mutation produced abnormalities in cardiac electrophysiology.

## Methods

### Generation of the Calm1 N54I mouse

A mouse carrying a CALM1 N54I mutation was generated using CRISPR/Cas9. The guide RNA targeting sequence contained a 20-bp sequence (GCAGGATATGATCAACGAAG) located in exon 3 of the Calm1 mouse gene, followed by the protospacer adjacent motif TGG. This sequence flanked by *Bbs*I sites was inserted into pX330, a vector expressing the guide RNA under control of the U6 promoter, and cas9 under a hybrid chicken β-actin promoter. The vector was injected alongside a single-stranded 192 base repair oligonucleotide into 147 × 0.5-day B6D2 mouse embryos. The repair oligo contained 91-bp homology arms, a AAC > ATC codon substitution, and a few additional third base mutations to prevent re-targeting of cas9 to the repaired DNA. Of 147 embryos injected, 105 survived and were transferred to 6 pseudo-pregnant females, thereby generating 23 pups. At weaning, genotyping was done by amplifying a 384-bp fragment, followed by sequencing. The PCR primers (forward, 5’ CAGTGAAGTGTGTGTCACAACAGTG 3’; reverse, 5’ ATTCTAATGTCAGGAGGCTGAGGC 3’) are located within the introns and therefore unique to CALM1. The sequencing primer (5’ TTGATTCTAAAGAGTCAAC 3’) is internal to the PCR fragment sequence. Three animals out of 23 were identified as having a correct mutant allele. All 3 lines were carried out by crossing to C57BL/6J female mice to demonstrate germline transmission. Line ‘6’ was further bred with the C57BL/6J mouse strain to dilute any possible off-target and strain effects. Heterozygote animals were bred with wild type to obtain heterozygote and wild type littermate controls.

### Calmodulin protein quantification

In-gel protein GluC digestion-Gel bands corresponding to the samples and calibration standards were excised and destained using a solution containing 50% aqueous acetonitrile (ACN) and 100 mM ammonium bicarbonate (ABC). The destained bands were cut into approximately 1-mm pieces, dehydrated with ACN, and dried in a SpeedVac concentrator (Thermo Fisher Scientific, Waltham, MA). The dried gel pieces were rehydrated in digestion buffer consisting of 10% aqueous ACN and 50 mM ABC. Proteins were subsequently reduced with dithiothreitol (DTT) and alkylated with iodoacetamide (IAM). The gel pieces were again dehydrated with ACN, dried in the SpeedVac, and rehydrated with the digestion buffer. Proteolytic digestion was performed with GluC at 37 °C overnight. Following digestion, the samples were centrifuged and the supernatants were recovered. The reactions were quenched by adding 6% formic acid, followed by centrifugation and collection of the supernatants. The remaining gel pieces were extracted with ACN, and the resulting supernatants were collected and combined with the previously recovered fractions. The pooled extracts were dried in the SpeedVac.

Nano-liquid chromatography nLC/MS/MS-The peptide digests were separated by nanoflow liquid chromatography (nLC) using an EASY-nLC II system (Thermo Fisher Scientific, Waltham, MA). Peptides were first loaded onto a 100 µm × 2 cm trap column for desalting (EASY-Column, catalog no. SC001; Thermo Fisher Scientific) and subsequently resolved on a 75 µm × 10 cm C18-AQ analytical column (catalog no. SC003; Thermo Fisher Scientific). The mobile phases consisted of 99.9% water and 0.1% formic acid (mobile phase A) and 99.9% ACN and 0.1% formic acid (mobile phase B). For the initial identification of peptides, separation was achieved using a 90-min linear gradient from 5% to 33% B at a flow rate of 300 nL/min. For subsequent peptide quantification, a 90-min linear gradient from 2% to 30% B was used. The column effluent was ionized online and detected using an LTQ Orbitrap Velos Mass Spectrometer (Thermo Fisher Scientific). During the initial peptide-identification analysis, precursor ions were measured in the Orbitrap at a resolution of 60,000 (at *m/z* 800), followed by data-dependent MS/MS analysis of the 10 most abundant precursor ions using collision-induced dissociation (CID) in the linear ion trap. For subsequent protein quantification, precursor-ion signals were acquired exclusively in the Orbitrap over targeted, narrow *m/z* ranges. The acquisition ranges were *m/z* 565–665 for the F90L mutation and *m/z* 845–945 for the N54I mutation.

Data Analysis-For peptide identification, the acquired Xcalibur RAW files were processed using Proteome Discoverer version 1.4 with the Sequest HT search engine (Thermo Fisher Scientific). Database searches were performed against a custom sequence database containing only human calmodulin and the corresponding mutant variants. Methionine oxidation and cysteine carbamidomethylation were included as variable modifications. For protein quantification, Xcalibur RAW files were processed using Xcalibur Quan Browser. Peak areas, expressed as areas under the curve (AUCs), were determined for the calibration standards and experimental samples. Calibration curves with coefficients of determination (R^2^) greater than 0.97 were achieved to calculate the corresponding peptide quantities in the experimental samples.

### Electrocardiographic recordings

All experiments were approved by the Vanderbilt University Institutional Animal Care and Use Committee. For telemetry recordings, electrocardiographic transmitters (DSI) were surgically implanted in mice under isoflurane anesthesia. The two recording electrodes were positioned in the left and right pectoral regions to approximate a lead I configuration. Following recovery, conscious mice were recorded while freely moving in their cages and during treadmill exercise. Treadmill exercise started at a treadmill speed of 100 m/minute and increased by 20 m/minute every two minutes until the mouse reached exhaustion, at which point it was returned to its cage.

Surface ECGs were recorded from anesthetized mice. Isoflurane concentration was titrated to the lowest level sufficient to maintain anesthesia, and four subcutaneous limb electrodes were used to record leads I and II. After establishment of a stable baseline heart rate, mice received intraperitoneal isoproterenol (3 mg/kg) and caffeine (100 or 120 mg/kg). ECG recordings were continued for 20 min following injection.

ECGs were analyzed by an investigator blinded to genotype, and ventricular and atrial arrhythmias were manually identified and annotated using LabChart.

### Cardiomyocyte Ca^2+^ measurements

Ventricular cardiomyocytes were isolated from wild-type and *Calm1*-N54I/+ littermates, incubated with 2 µM Fura-2 AM for 7 minutes, and washed 2 × 10 minutes with normal Tyrode (NT) solution containing 250 µM probenecid. The composition of NT used for Fura-2 loading and washing was (in mM): 134 NaCl, 5.4 KCl, 1.2 CaCl^2^, 1 MgCl^2^, 10 glucose, and 10 HEPES, pH adjusted to 7.4 with NaOH. After Fura-2 loading, all experiments were conducted in NT solution containing a Ca^2+^concentration of 2 mM. Fura-2 loaded myocytes were recorded at 3 Hz field stimulation for 20 seconds after application of superfused NT solution containing 1 µM isoproterenol with drug, followed by no electrical stimulation for 40 seconds to record spontaneous Ca^2+^release events. After that, myocytes were perfused with 10 mM caffeine in NT solution for 5 seconds to estimate total SR Ca^2+^content. Fura-2 was measured using a dual-beam excitation fluorescence photometry setup (IonOptix Corp.) and analyzed using commercially available data analysis software (IonWizard, IonOptix, Milton, MA). All experiments were conducted at room temperature.

### Statistical analysis

Statistical analyses were performed using GraphPad Prism or R for Windows. Statistical tests and sample size are indicated in the corresponding figure legends. A two-sided P value <0.05 was considered statistically significant.

## Results

### Generation of a heterozygous Calm1 N54I mouse

We used CRISPR/Cas9-mediated genome editing to generate a mouse carrying the N54I mutation in *Calm1*. Targeting of exon 3 introduced an AAC-to-ATC nucleotide substitution, resulting in replacement of asparagine with isoleucine at residue 54 of mouse CaM (Figure 1A–C). Three founder lines carrying the intended mutation were identified and bred to establish germline transmission. One line was subsequently backcrossed onto the C57BL/6J background to reduce potential effects of off-target editing and genetic background.

**Figure 1.**
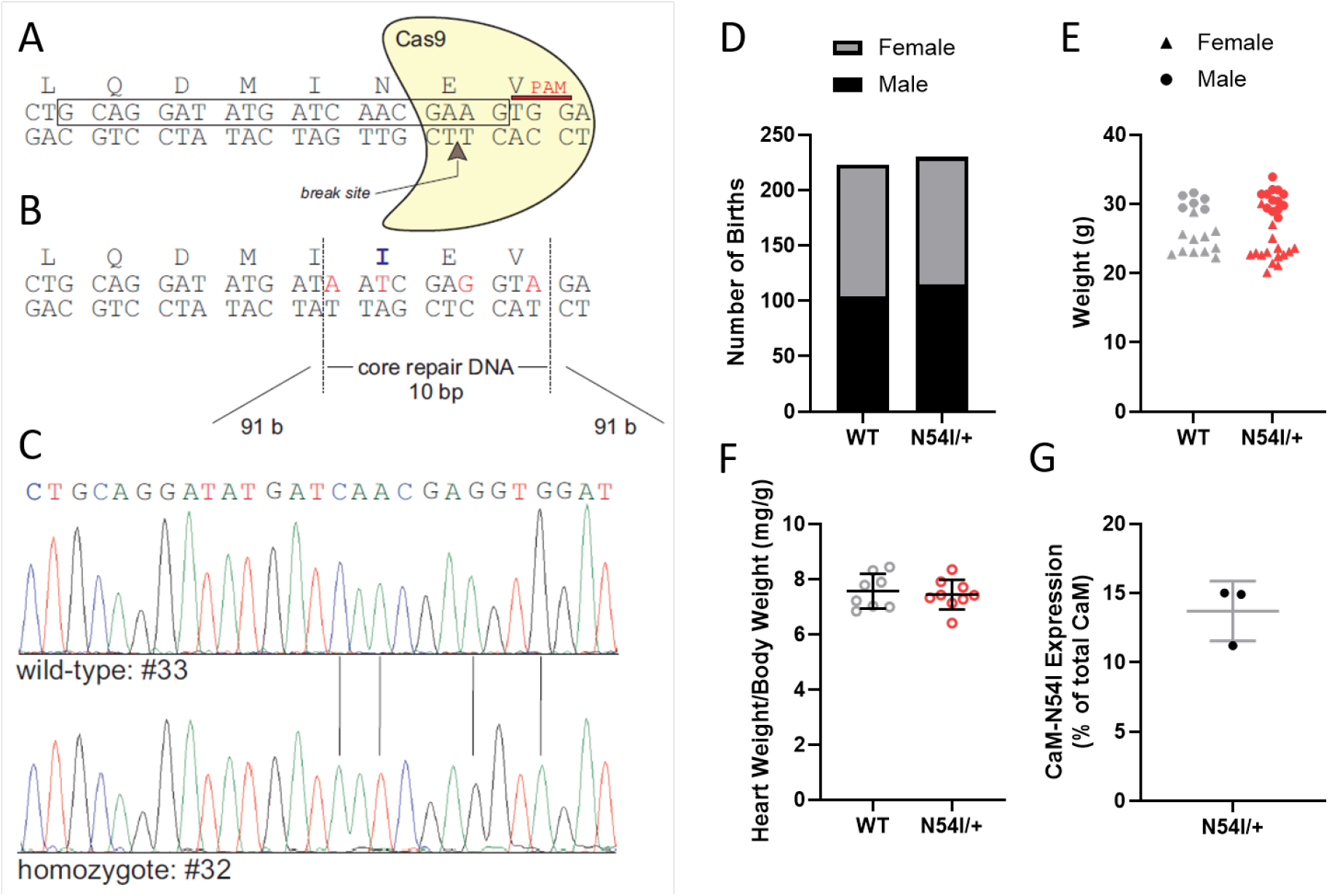
Generation and characterization of calmodulin (CaM) N54I/+ mouse. A) Guide RNA targeting sequence for gRNA (boxed region) and the resulting break site. B) The mRNA and protein sequence flanking N54, resulting in an isoleucine. C) DNA sequencing results for selected wild-type (WT) and CaM-N54I lines. D) Number of births by genotype and sex. E) Body weight at 18-24 weeks of age by genotype and sex. F) Heart to body weight ratios for wild type (WT) and N54I/+ littermates. G) CaM-N54I protein expression in hearts isolated from three CaM-N54I/+ mice.

Subsequent generations were bred by crossing wild type (WT) and heterozygous CaM-N54I/+ mice. Breeding produced litters of normal size and genotype distributions consistent with the expected Mendelian ratio (Figure 1D). CaM-N54I/+ mice exhibited normal growth (Figure 1E) and behavior. Seven out of 230 heterozygotes died before 12 weeks of age of unknown cause (P = 0.015, Fisher’s exact test vs WT littermates). Heart weight to body weight ratio was comparable between *Calm1*-N54I/+ mice and wild-type littermates (Figure 1F).

Because N54I is expressed from only one of the six alleles encoding CaM, we next quantified mutant CaM in the heart. Mutant CaM accounted for 13.7% of total cardiac CaM in *Calm1*-N54I/+ mice (Figure 1G). This value is consistent with approximately equivalent expression of the mutant allele relative to the other five wild-type *Calm* alleles.

### Calm1 N54I/+ mice exhibit normal baseline cardiac electrophysiology

We next determined whether the N54I mutation altered baseline cardiac electrophysiology. Surface electrocardiograms (ECGs) were recorded from anesthetized mice under conditions in which isoflurane concentration was minimized to reduce effects on cardiac electrophysiology. All ECGs were analyzed by an investigator blinded to genotype. At baseline, *Calm1*-N54I/+ mice exhibited normal ECG parameters and morphology (Figure 2A&B). Heart rates were comparable to wild-type littermates (Figure 2C), and P duration, PR interval, QRS width, and QT interval were also not significantly different between genotypes (Figure 2D–G). These findings indicate that the N54I mutation does not produce a major baseline abnormality in atrioventricular conduction, ventricular depolarization, or ventricular repolarization. In particular, the absence of QT prolongation is consistent with the CPVT rather than LQTS phenotype associated with N54I in humans, and with the normal calcium binding properties of recombinant CaM-N54I. The normal baseline ECG also suggests that the mutation does not produce a sufficiently large global disturbance of sarcolemmal ion-channel function to alter conventional surface ECG parameters.

**Figure 2.**
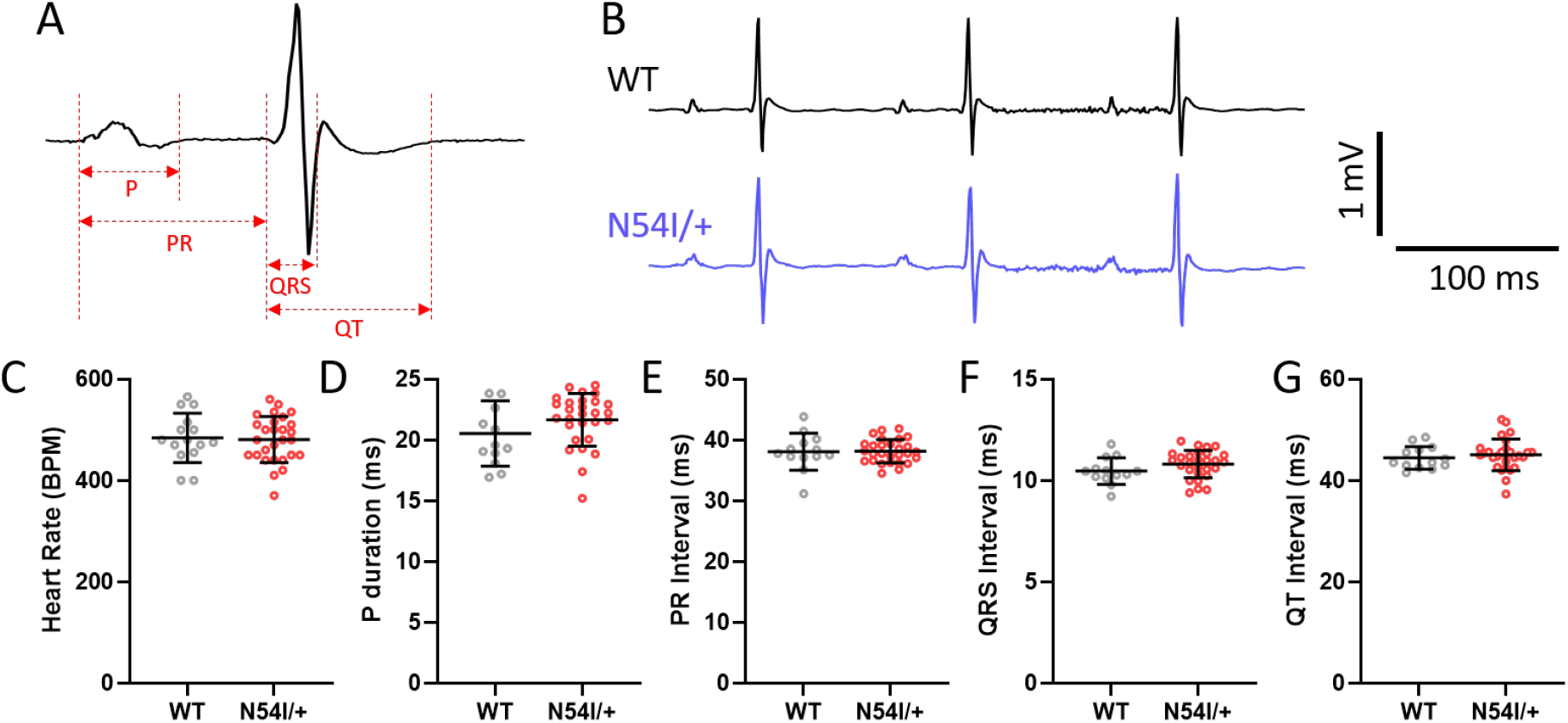
Baseline electrocardiogram (ECG) parameters from anesthetized wild type (WT) and CaM-N54I/+ mice. A) Representative lead I ECG showing P duration (P), PR interval, QRS interval, and QT interval. B) ECG recordings from a WT and CaM-N54I/+ mouse showing normal waveform morphology. C) Heart rate, D) P duration, E) PR interval, F) QRS width, and G) QT interval. P > 0.05 by Student’s t-test for all parameters. Data shown as mean±SD. N = 13 (WT) and 28 (N54/+) mice.

### Calm1 N54I/+ cardiomyocytes exhibit hyperactive spontaneous Ca^2+^ release

To determine whether the N54I mutation produced a cellular Ca^2+^ phenotype, ventricular cardiomyocytes were isolated from *Calm1*-N54I/+ mice and wild-type littermates and loaded with Fura-2 AM. Paced Ca^2+^ transient properties were unchanged (Figure 3A–F). SR calcium load was assessed by application of 10 mM caffeine. There were not any appreciable differences in SR Ca^2+^ in cardiomyocytes from CaM-N54I/+ mice (Figure 3G–H). To evaluate the impact of mutant CaM on RyR2-mediated Ca^2+^ release, cells were stimulated at 3 Hz followed by a 40-second period of cessation of pacing, during which spontaneous Ca^2+^ release events with or without triggered beats could be recorded (Figure 3B). Cardiomyocytes from CaM-N54I/+ had significantly more spontaneous calcium release events and triggered beats (Figure 3I– J).

**Figure 3.**
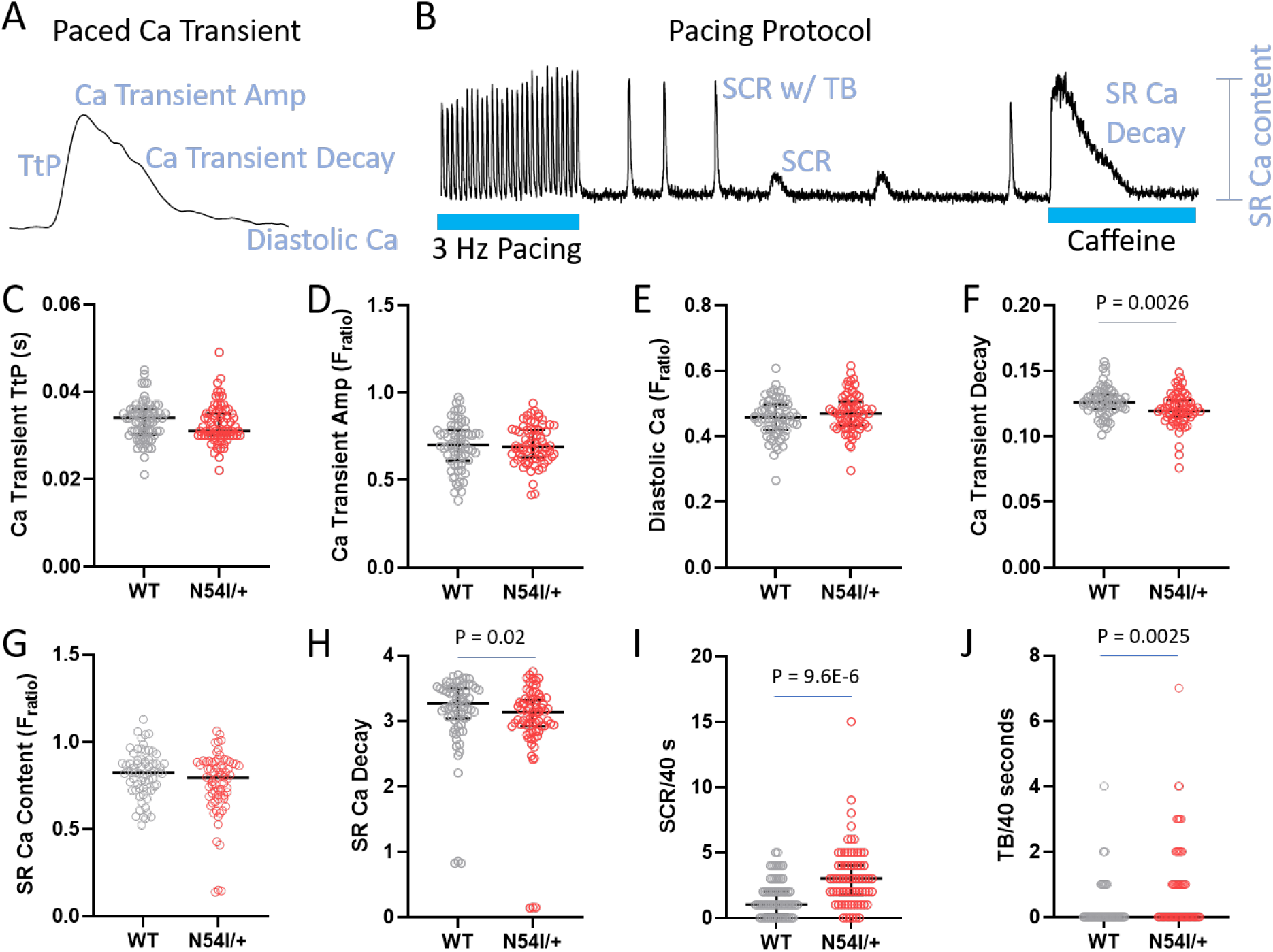
Cardiomyocytes from CaM-N54I/+ mice have increased spontaneous calcium release. A) Representative calcium (Fura-2) recording from a paced cardiomyocyte, illustrating various parameters. B) Pacing protocol used to elicit spontaneous calcium release (SCR) events and triggered beats (TB) following cessation of pacing. SR = sarcoplasmic reticulum. C-F) Paced calcium transient parameters from 3 Hz pacing (see also panel A). G) SR calcium content induced by application of 10 mM caffeine. H) SR calcium decay constant. I) Fractional calcium release relative to the SR content. J) Spontaneous calcium release events over the 40-second cessation of pacing. N = 68 cells (WT) and 70 cells (N54I/+ from 3 mice per group. Data compared using a mixed-effects hierarchical clustering model.

### Calm1 N54I/+ mice develop catecholamine-induced ventricular arrhythmias

To determine whether the cellular Ca^2+^ phenotype translated into ventricular arrhythmias in vivo we performed a catecholamine challenge. Anesthetized mice were challenged with intraperitoneal administration of isoproterenol (3 mg/kg) and caffeine (100 or 120 mg/kg). Following catecholamine challenge, heart rate increased (Figure 4A) to a similar degree in CaM-N54I/+ mice as WT (Figure 4B). Catecholamine challenge produced varying degrees of ECG changes (Figure 4C–D). We observed hallmark ventricular arrhythmias consistent with a CPVT phenotype, chiefly premature ventricular contractions (PVCs) and bigeminy, but also ventricular tachycardia (Figure 4C). CaM-N54I/+ mice had significantly greater rates of ventricular ectopy compared with WT mice (Figure 4E). CaM-N54I/+ mice also had more severe arrhythmias as assessed by an ordinal scoring system, with many of them progressing to non-sustained ventricular tachycardia (Figure 4F). One WT mouse had a single couplet.

**Figure 4.**
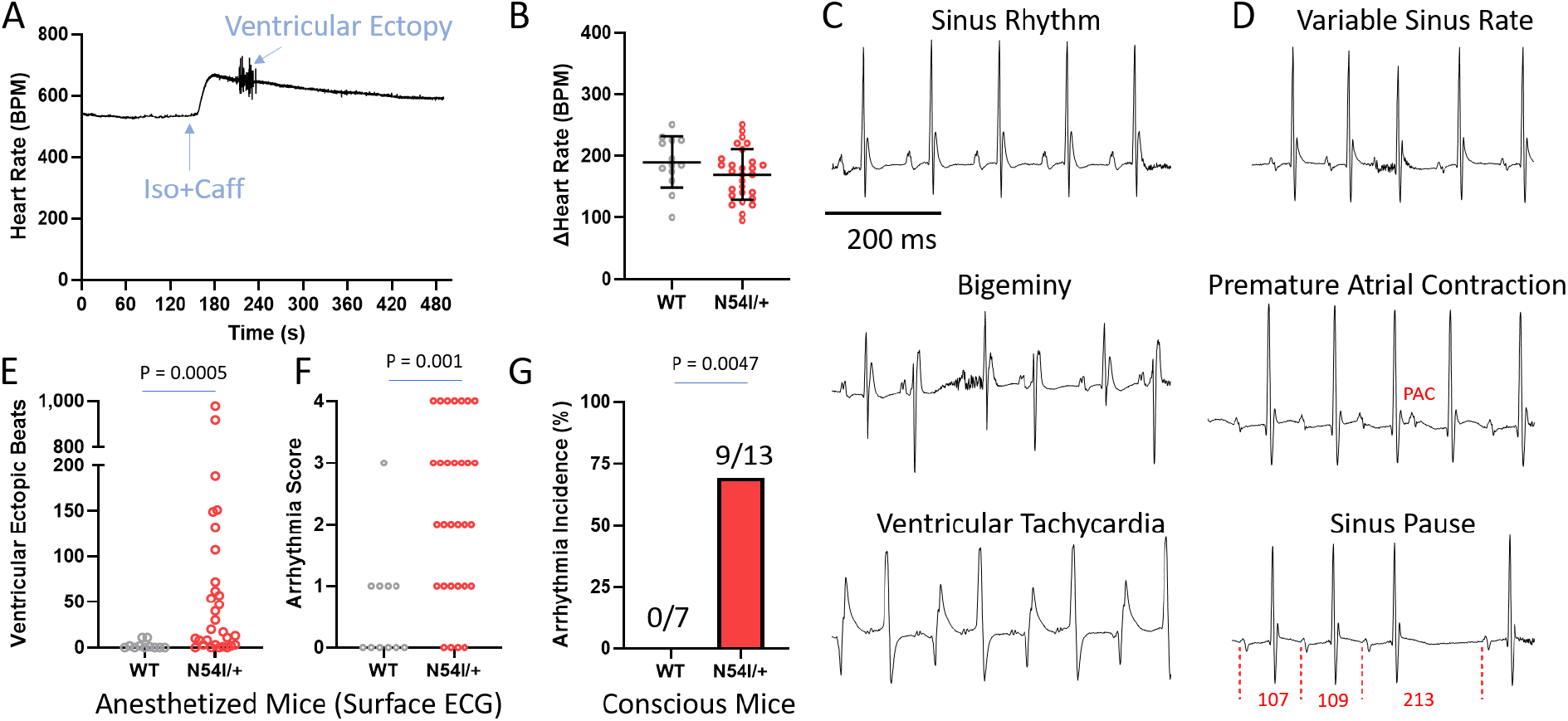
Arrhythmia phenotypes in CaM-N54I/+ mice. A) Heart rate recording in an anesthetized CaM-N54I/+ mouse showing an increase in response to intraperitoneal administration of isoproterenol and caffeine. Heart rate response became variable during a period of ventricular ectopy. B) Heart rate response to catecholamine challenge. C) Representative electrocardiogram recordings showing normal sinus rhythm (top), bigeminy (middle), and bidirectional ventricular tachycardia (bottom). D) Representative electrocardiogram recordings of atrial sinus node dysfunction. PAC = premature atrial contraction. The numbers in the bottom trace indicate the interval (in ms) between P waves. E) Total number of ventricular ectopic beats. F) Arrhythmia severity score. 0 = no ventricular ectopy, 1 = isolated premature ventricular contractions, 2 = bigeminy, 3 = couplets, 4 = non-sustained ventricular tachycardia. N = 11 (WT) and 30 (N54I/+) mice in panels E and F. Data compared using Mann-Whitney U test. G) Ventricular arrhythmia incidence measured by telemetry in conscious mice undergoing treadmill exercise. P = 0.0047 by Fisher’s exact test.

To determine whether ventricular arrhythmias could also be provoked under physiological conditions in conscious animals, mice were implanted with ECG telemetry and subjected to treadmill exercise. Exercise provoked ventricular arrhythmias in 9 of 13 *Calm1*-N54I/+ mice (Figure 4G). We also noted the presence of PVCs in conscious mice at baseline/rest (data not shown). The stress-dependent appearance of ventricular ectopy and tachycardia, together with the absence of a major baseline ECG abnormality (such as prolonged QT interval), is consistent with a CPVT phenotype.

### Calm1 N54I/+ mice exhibit atrial and sinoatrial abnormalities

In addition to ventricular arrhythmias, ECG recordings revealed abnormalities in atrial electrical activity in many of our CaM-N54I/+ mice. These abnormalities included variable atrial conduction, premature atrial contractions, ectopic and occasionally multifocal P waves, intermittent dropped beats consistent with conduction block, and beat-to-beat variability in atrial activity (Figure 3D). Abnormal sinoatrial and atrial electrophysiology has also been described in other CPVT models. In *Casq2*−/− mice, loss of calsequestrin produces sinus node dysfunction, altered atrial Ca^2+^ cycling, conduction abnormalities, and increased susceptibility to atrial arrhythmias.^14^ These findings suggest that abnormal SR Ca^2+^ handling can influence pacemaker and atrial electrophysiology in addition to ventricular arrhythmogenesis.

## Discussion

The principal finding of this study is that expression of the N54I calmodulin mutation from a single *Calm1* allele is sufficient to produce a stress-induced ventricular arrhythmia phenotype in vivo. Our data establish that the N54I mutation produces a bona fide CPVT phenotype. This finding is particularly important in the context of the unusual genetic organization of the *CALM* genes, where a heterozygous mutation in one *CALM* gene is predicted to contribute only a small fraction of the total CaM pool.^7^ Our data confirmed experimentally (**Fig. 1**) that mutant CaM-N54I represented only 13.7% of total cardiac CaM. Nevertheless, this relatively small fraction of mutant CaM was sufficient to produce a clear arrhythmia phenotype.

Our findings therefore provide *in vivo* evidence for the functional dominance of CPVT-associated CaM mutations. This conclusion extends previous *in vitro* work from our group demonstrating that CPVT-associated CaM mutants can exert dominant effects on RyR2 because they exhibit increased binding affinity to RyR2 compared to WT-CaM, but unlike WT-CaM, do not inhibit RyR2 channel activity.^12^ As a result, N54I and other CPVT-associated CaM variants increased spontaneous Ca^2+^ waves and sparks in ventricular cardiomyocytes and increased RyR2 channel open probability. Importantly, a mixture containing only 1 part mutant CaM to 8 parts wild-type CaM was sufficient to promote spontaneous Ca^2+^ release.^12^

The present findings also extend the original clinical discovery of the N54I mutation. The human N54I variant was initially identified in a family with dominantly inherited CPVT-like ventricular arrhythmias and was subsequently shown to segregate with disease.^4^ Although the clinical genetic evidence implicated the variant as pathogenic, the present mouse model provides an independent line of evidence.

The cellular mechanism is consistent with a primary defect in SR Ca^2+^ release. Our previous in vitro work demonstrated that recombinant CPVT-associated CaM mutants can directly alter RyR2 regulation, increasing channel activity and promoting SCR.^12^ In ventricular cardiomyocytes isolated from N54I/+ mice, we observed a similar increase in SCR (**Fig. 3**). Such spontaneous diastolic Ca^2+^ release provides a substrate for triggered activity through activation of the electrogenic Na^+^/Ca^2+^ exchanger. At the tissue level, studies indicate that spontaneous Ca^2+^ release and DADs in ventricular myocardium near the Purkinje–myocardial junction can generate full action potentials in adjacent Purkinje fibers, providing a mechanism by which cellular Ca^2+^ abnormalities can be translated into premature ventricular beats.^9^

An important feature of the present model is the absence of a major baseline ECG abnormality. Heart rate, PR interval, QRS duration, and QT interval were comparable between N54I/+ and wild-type littermates (**Fig. 2**). This finding is particularly relevant because CaM mutations can produce clinically distinct phenotypes, including CPVT, LQTS and overlap syndromes.^15^ The normal QT interval in N54I/+ mice is therefore consistent with the CPVT phenotype previously associated with N54I rather than with a generalized repolarization abnormality.

Although the primary phenotype of N54I is ventricular arrhythmia, several N54I/+ animals exhibited premature atrial contractions, ectopic P waves, variable conduction, dropped beats, and beat-to-beat variability. Similar abnormalities have been reported in other CPVT models. *Casq2*−/− mice have sinoatrial node dysfunction, conduction abnormalities, and atrial arrhythmias, demonstrating that perturbation of intracellular Ca^2+^ handling can affect pacemaker and atrial electrophysiology.^14^ The lack of baseline bradycardia in N54I/+ mice distinguishes this model from *Casq2*−/− mice. In the latter, increased diastolic Ca^2+^ and abnormal Ca^2+^ release within sinoatrial node cells contribute to pauses, unstable pacemaking, and reduced heart rate.^14^ The N54I/+ phenotype therefore appears to produce a more selective alteration in atrial electrical stability without the pronounced suppression of sinus rate, though it should be noted that patients carrying the N54I mutation reportedly had bradycardia.^4^ The mechanistic basis for this distinction remains to be determined.

Our findings should also be considered in relation to the previously described *Calm1* N98S mouse model. N98S is another CPVT-associated CaM mutation and was subsequently modeled using a heterozygous knock-in strategy. However, that model produced a complex arrhythmia phenotype including LQTS, QRS widening, CPTV, and sinus bradycardia.^16^ Unlike in the N54I model presented here, mutant N98S protein is enriched in heart tissue due to a decrease in N98S-CALM protein degradation.^17^ Furthermore, unlike N54I, N98S has modestly reduced Ca-binding affinity,^12^ which may contribute to prolonged repolarization and sex-differences observed in N98S mouse models.^18^

There are several limitations to the present study. First, CaM regulates numerous cardiac proteins, including CaV1.2 and other ion channels, and additional effects of N54I cannot be excluded.^7^ Second, the present study does not directly determine whether mutant and wild-type CaM occupy identical molecular pools or whether N54I preferentially associates with specific cardiac targets. Third, the mechanisms responsible for the atrial and sinoatrial abnormalities remain unresolved.

Despite these limitations, the present study establishes an important biological principle: a pathogenic CaM mutation expressed from a single allele is sufficient to produce a clinically relevant arrhythmia phenotype in vivo. The observation that mutant CaM constitutes only 13.7% of total cardiac CaM yet produces a robust stress-induced ventricular arrhythmia phenotype provides experimental support for the functional dominance of CPVT-associated CaM mutations. More broadly, these findings underscore the critical role of this signaling protein, and how it can produce profound abnormalities in cardiac Ca^2+^ handling and electrical stability.

## NON-STANDARD ABBREVIATIONS AND ACRONYMS

CaM: Calmodulin
Casq2: calsequestrin, cardiac isoform
CaMKII: Ca/CaM kinase II
CPVT: Catecholaminergic Polymorphic Ventricular Tachycardia
LQTS: Long QT Syndrome
RyR2: Ryanodine receptor Ca release channel
SR: Sarcoplasmic Reticulum

